# The conditional defector strategies can violate the most crucial supporting mechanisms of cooperation

**DOI:** 10.1101/2022.06.07.495117

**Authors:** Ahmed M. Ibrahim

## Abstract

Cooperation is essential for all domains of life. Ironically, it is intrinsically vulnerable to exploitation by cheats. Hence, there is an explanatory necessity that triggers a lot of evolutionary biologists to search for mechanisms that could support cooperation. In general, cooperation can emerge and be maintained when cooperators are sufficiently interacting with themself to provide a kind of assortment and reciprocity. One of the most crucial and common mechanisms to achieve that task are kin selection, spatial structure, and enforcement (punishment). Here I used agent-based simulation models to investigate these pivotal mechanisms against conditional defector strategies and concluded it could easily violate all of them and take over the population. This surprising outcome may cue us to rethink the evolution of cooperation as it illustrates that maintaining cooperation may be more difficult than previously thought. Moreover, besides the theoretical findings, there are empirical applications such as invading the cooperator population of pathogens by genetically engineered conditional defectors, which could be a potential therapy for many incurable diseases.

## 1. Introduction

A long-standing puzzle in evolutionary theory is how cooperative behavior can evolve and persist within the selfish natural world. Once cooperation exists, it is always vulnerable to exploitation by defective free riders who adopt selfish strategies for reaping the highest possible profit without paying the share. Thus, invest most of their energy in reproduction. Therefore, cheaters could out-compete the cooperators and take over the population. Since Haldane pointed out that there was no general principle to solve this problem (Ferriere & Legendre 2013). Many partial mechanisms have been suggested, Such as kin selection, group selection, reciprocity, policing, spatial structure, sanction, reward, and punishment (West 2007; Nowak 2006; Cremer et al. 2019). Darwin himself had suggested some core concepts of these mechanisms. In this paper, I introduced conditional defector strategies that violate Kin selection, punishment, and spatial structure mechanisms.

### 1.1. What are conditional defector strategies?

From a conceptual viewpoint, a conditional defector strategy may be any cheating strategy that could somehow cooperate. In other words, it is a cheater who pays additional costs or wastes a portion of the profit to survive. So, it is not obligated to defect in all behaviors or at all times. However, it may cooperate in some behaviors, now and then; it may cooperate with some agents, etc. So they are not pure defectors

### 1.2. Some forms of conditional defector strategies

#### 1.2.1. Cooperate for the spread

Dispersal is beneficial because it decreases Kin competition, sustains the resources, and outset colonization. Without dispersal, the fate of all populations is extinction, (Bonte & Dahirel 2017). Hence, an intermediate dispersal rate of cooperators is essential for cooperation maintenance, (Parvinen 2011; Waite et al. 2015). Nevertheless, dispersal is a costly behavior that increases the mortality of dispersers or decreases their fecundity, (Bonte et al.2012; Lion & Baalen 2008). Therefore, some studies focused on the joint evolution of dispersal and cooperation, (Parvinen 2013). Or the correlation between cooperation, dispersal rate, and dispersal cost, (Galliard et al. 2005). Their findings asserted that the low dispersal cost selects against cooperation. Thus, if cheaters can reduce their dispersal costs, they may turn the game against cooperators. Usually, cheats are not good migrators because dispersal itself is a cooperative behavior. The migrators leave their suitable habitats to other unknown environments and face dangerous predators to colonize a new patch. Such behavior is costly for the migrators. Nevertheless, its benefits are also gained by non-migrators because it decreases kin competition. From such a point of view, dispersal is considered cooperative behavior that naturally does not expect to be abundant in cheats. Hence, cheaters go extinct rapidly with the depletion of local patches they dominated without global prevalence like cooperators; this might be the fundamental problem of cheaters. However, the probable solution is adopting a conditional defection strategy wherein free-riders would cooperate only for the spread. The actors of this selfish strategy would have a high dispersal rate with the lowest possible cost because they share migration costs. Thus, the exploitation rate of public goods and interactions among defectors and cooperators will increase. In other words, the conditional defectors can exclusively cooperate for all collective behaviors related to migration (coalition dispersal) but defect otherwise. Therefore, these selfish successful migrators can convert the structured meta-population into a well-mixed game and violate the spatial structure mechanism.

(Ridley 2012) is considered a piece of empirical evidence for the assumption that individuals reduce their dispersal costs by sharing it. Thereby they can achieve successful migrations. Also, in metastases cancer, migrating in groups (coalition dispersal) raises the efficacy up to 50-fold more than individual dispersal, (Tissot et al. 2019; Kümmerli et al. 2009).

#### 1.2.2. Pay for the escape

Conditional defectors can pay some of their wealth or waste some profits to escape punishment by producing substances to mislead punishers. Or possession of the significant tag that marks cooperators. Similarly, by reducing their payoff to be more similar and familiar to cooperators. If cheaters reduced the benefits, it might be hard to have been noticed by a quorum-sensing system or other defense mechanisms. It is considered a kind of imitation or tag-based decision that prevents cooperators from detecting and punishing the defectors.

Anyway, those cheaters pay a cost to escape sanction or reduce the accuracy of the monitoring/punishment system. Therefore, they can merge with cooperator populations accordingly, violating punishment and kin-selection mechanisms.

## 2. Methods

I used two agent-based simulation models to investigate the concepts of “cooperate for the spread” and “pay for the escape” both of them are Net logo models created by Dr. Susan Hanisch.

Afterward, I modified the first model to represent the concept of sharing the dispersal costs. I used the second model without modifications. But instead, I assigned definite values of some parameters that highlight the pay for the escape strategy.

### 2.1. First model

The original model was entitled “Evolution and patchy resource”, (Hanisch 2017a). She developed the model in the first place for educational purposes. The model illustrates concepts of cooperators-cheaters competition, natural selection, spatial structure mechanism, multilevel selection, and founder effect.

#### 2.1.1. Changeable variables

- Distance-resource-areas: the distance between the centers of the resource areas.
- Size-resource-areas: the size of resource areas as a radius in the number of patches.
- Living costs: the costs that each agent has to deduct from energy per iteration for basic survival.
- Mutation rate: The probability accordingly offspring agents have different traits than their parents.
- Evolution: the ability of agents to produce offspring.

#### 2.1.2. Constant variables

- The number of patches is 112 * 112 patches.
- Carrying capacity per patch: Resource = 10, Agents = 1
- The growth rate of the resource = 0.2
- The resources on a patch regrow by a logistic growth function up to the carrying capacity: New resource level = current resource level + (Growth-Rate * current resource level) * (1 - (Current resource level / carrying capacity).
- The cost for producing offspring is 10 subtracted units of energy.
- The initial level of energy of agents is set at living costs.

#### 2.1.3. Role of randomness

- Agents are distributed randomly in resource areas at the beginning of a simulation.
- Sustainable behavior is distributed randomly with a probability of Percent-Sustainables among the initial agent population.
- The order in which agents move and harvest within one iteration is random.
- Agents move to a randomly selected patch if several patches fulfill the objectives.
- The order in which agents produce offspring within one iteration is random.
- Agents reproduce offspring with a probability of (0.0005 * Energy).
- Agents place offspring on a randomly selected unoccupied neighboring patch.
- Offspring mutate with a potentiality of Mutation-rate.

#### 2.1.4. Model Processes

In each iteration, each agent moves around in random order. There are three likelihoods:

- If there are no unoccupied patches in 2 patch radius, they stay on the current patch.
- If there are unoccupied patches with resources amounting to more than living costs, the agents move to them.
- If the resource amount was less than the living costs, the agents move randomly to other unoccupied patches.

The agents harvest the resources from separated patches to gain energy for metabolism and proliferation. If the energy level of any agent falls to zero, it dies. The cooperator type harvests half of the resource, while the greedy type consumes 0.99 %.

The living costs are deducted from the energy amount of the agent constantly everywhere all the time. This process occurs regardless of whether an agent moves within the patch, between the patches, or even not moving. Therefore, the model doesn’t consider dispersal cost explicitly.

If there is an unoccupied neighbor patch, the agent can reproduce in probability 0.0005 of his energy and place the offspring on the unoccupied neighbor patch, then transfer 10 units of the energy to his offspring.

Resources regrow only on resource patches. If the resource amount is more than or equal to 0.1. Then it regrows. If the resource is less than 0.1. it is set to be 0.1.

#### 2.1.5. Output diagrams and monitors

- The average energy of agents: average energy levels of sustainable and greedy agents, resulting from resource harvest, minus living costs and reproduction.
- Trait frequencies: The relative frequencies of sustainable and greedy agents in the total population, resulting from mutations, different reproduction rates, and death.
- Agent Population: The absolute number of the total population size, resulting from reproduction and death.

#### 2.1.6. Modifications

In the first modification, I added a different type of cost that agents only incur when they disperse from one patch to another (in-between the patches). It is the slider entitled “dispersal-costs.”

In the second modification, I added another sharing dispersal costs tool to reduce them by dividing their value by the sum number of included agents (flock-mates) in the identified range from the same type. It is the slider entitled “group-dispersal-range.” which is the flock mate’s areas as a radius in the number of patches. So, changing the value of the group dispersal range will change the area around every agent. Accordingly, the number of its flock mates who share the dispersal costs also changes.

The group dispersal range is variable and not exclusively for greedy agents but applies to all agents. So, it represents the case of the wild-type of cooperators who also can cooperate for the spread. The group dispersal range also does not only target the agents in between patches. But it counts the agents inside and outside the patches. For example, once an agent starts its dispersion with a determined range containing 10 agents, 4 from another type, 3 non-dispersal agents from the same type that existed inside a patch, and 3 dispersal agents from the same type outside the patches. The dispersal costs for this agent will be divided by 6. This case may represent in the real world via public good of diffusible stuff or similar techniques.

Cheaters can arise within cooperator patches by mutation or immigration. Therefore, to investigate the efficacy of migration, the mutation rate value should be 0 to cancel its effect in the meta-population dynamics.

### 2.2. Second model

The model entitled “Evolution, resources, monitoring, and punishment.” (Hanisch 2017b), is a simulation of a population with four types of agents competing for the same resource. It demonstrates many concepts like kin selection, cooperation, selfishness, public good, monitoring, punishment, sharing the costs, positive /negative frequency-dependent selection, and multilevel selection. The four agent colors and types: 1) Red: greedy, non-punishing. 2) Orange: greedy, punishing. 3) Turquoise: sustainable, non-punishing. 4) Green: sustainable, punishing.

Punishing agents can perceive other agents in their environment to some degree called (perception accuracy) and react to their behavior. There are three kinds of punishment: Punishers can kill agents with greedy harvesting behavior, stop them from harvesting in the next iteration (I selected this kind), or have them pay a penalty fee to their neighbors.

Agents have a cost (energy) to pay for, both detection and punishment, so this behavior is altruistic. Punisher agents of one type share punishment cost equally.

#### 2.2.1. Changeable variables

- Death rate: The probability accordingly agents die independent of their energy level.
- Carrying capacity: the maximum amount of resource units on a patch from 1-to 100.
- Growth rate: the rate at which resources on patches regrow. The maximum sustainable yield is calculated based on carrying capacity and growth rate.
- Harvest-sustainable: the number of resource units harvested by sustainable agents.
- Harvest-greedy: the number of harvested resource units by sustainable agents.
- Perception-accuracy: the probability with which punishing agents notice greedy agents.
- Costs-perception: the costs in units of energy, punishing agents have to pay for perceiving other agents.
- Costs-punishment: the costs as units of energy that punishing agents have to pay in each iteration for punishing other agents. All punishing agents of an agent divide the costs of punishment.
- Punishment: the kinds of punishing behavior that punishing agents perform.
- Fine: if the kind of punishment is “pay fine” the fine in energy units that punished agents have to pay (shared between all their neighbors).
- Living costs and Mutation-rate: see the first model.

#### 2.2.2. Constant variables

- The number of patches: There are 60*60 patches in the world.
- The initial energy level of agents is set at living costs + 1.
- The initial amount of resource units on a patch is set to Carrying-capacity.
- The resources on a patch regrow: see the first model.

#### 2.2.3. Role of randomness

* In addition to items in the first model.

- Agents take on their traits (harvest preference and ability to notice and punish) randomly based on the probability of Percent-sustainable and Percent-punishers.
- The order in which punishing agents notice greedy agents within one iteration is random.
- Greedy agents get noticed by punishing agents with a probability of Perception-accuracy.
- The order in which detected greedy agents get punished within one iteration is random.
- Agents produce offspring with a probability of (0.001 * Energy).
- Agents die with a probability of Death-rate.

#### 2.2.4. Model Processes

In each iteration, each agent attempts to harvest resources from the patches it is on and the eight neighboring patches until the harvest preference level is reached, except for the punished agent with the sanction (suspend harvest once), its harvest amount = 0 in the current iteration. If the amount of resources available is lower than the amount that the unpunished agent attempts to harvest. Then the agent moves to a neighboring unoccupied patch with the most resources after losing one energy unit as a move cost.

Punishers pay the costs of perception for sense greedy agents. The greedy neighbors have been noticed with the probability of perception accuracy. The agent lost an amount of energy as living costs. The agent dies with the probability of death rate or if the energy level falls to zero.

If there is an unoccupied neighbor patch, the agent can reproduce in probability 0.001 of its energy and place the offspring on the unoccupied neighbor patch, then transfer half of its energy to its offspring that mutate with the probability of mutation rate.

Resources regrow on all patches. If the resource amount is more than or equal to 0.1. Then it regrows. If the resource is less than 0.1. it is set to be 0.1.

#### 2.2.5. Output diagrams and monitors

- Populations (% of carrying capacity): The state of the resource and the agent population in the world as a percentage of total carrying capacity; resulting from resource harvesting behavior and resource regrowth, agent reproduction, and death.
- Average harvest per iteration: The average harvested amounts of agents per iteration by trait, resulting from harvested resource units, minus costs for monitoring and punishing (for punishing agents), minus fines (for punished agents in case of punishment “Pay fine”)
- The average energy of agents and Trait frequencies: see the first model.

#### 2.2.6. How does the model represent a conditional defector strategy?

The model goal is to highlight the role of kin selection and punishment mechanisms in supporting cooperation evolution against cheats. I didn’t need to modify the model but just thought about what the conditional defector should do to upside down the game? The answer was to pay for the escape.

For instance, if the standard Harvest-greedy of a cheater (greedy, non-punishing) was 13 and the Perception-accuracy of its actual punishers was 75%. Now suppose this cheater faces troubles, and it cannot dominate. But if it gives up some of its profit to become 12, to escape punishment, and to reduce the Perception-accuracy to 60%, it could dominate and take over the population.

The conditional cheater can pay something and reduce its profit to escape punishment by reducing Perception-accuracy if there is a positive correlation between the values of these two variables, which allow the cheater’s dominance. Therefore, this model is proper if it can support/deny such correlation.

## 3. Results

All experiments I carried out via a built-in tool in the Net logo called Behavior Space. And all data analyses I carried out via a Python library called Glueviz and excel.

### 3.1. The experiments of the first model

The default values of the variables: Mutation rate = 0, (to investigate only the effect of dispersal). Dispersal-costs = 8, (high value). The agent’s shape is Bacteria. Size-Resource-Areas = 4, (Relatively small). Living-costs = 1. Percent-Sustainables = 90%, (most of the population consists of cooperators in the beginning). Number-Agents = 80, (started number). Distance-Resource-Areas = 20, (Relatively far). Evolution switch is true, (natural selection is working). Group-dispersal-range = 0, 30, 50, 70,100,150, and 200.

They are 63 runs of 7 experiments. 15 repeated runs for group dispersal range = 0, and 8 repeated runs for each other value. Approximately all runs with group dispersal range= 0, finished in favor of cooperators and the extinction of cheaters as expected that cheaters cannot sustain their patches and cannot arrange successful migrations to other patches due to the high dispersal costs.

This situation significantly changed in the rest runs of group dispersal ranging from 30 to 200 where cheaters can share the dispersal costs. Consequently, all these runs finished in favor of cheaters, and all cooperators were extinct. Fig 1. Also, cheaters in these runs outcompete cooperators quickly with a fewer number of steps as long as the group dispersal range increases from 30 to 70. Then the average of steps is somewhat convergent for the group dispersal range from 70 to 200. Fig. 2. And Fig. 3.

**Figure.1:**
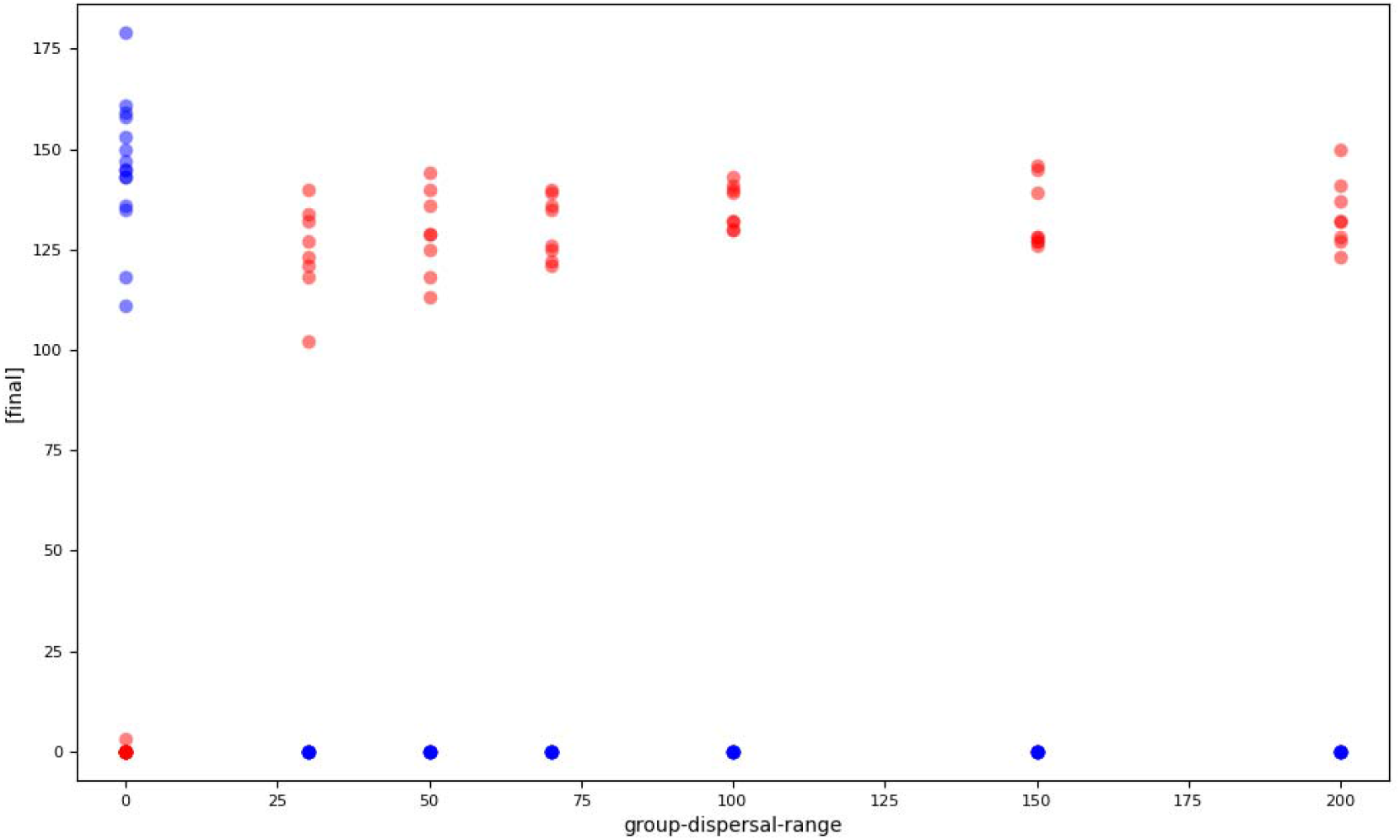
Cheaters (red) and cooperators (blue). Cheaters could thrive only when they start to share the dispersal costs to some degree. When group dispersal range = 0. Each cheater pays the dispersal costs by itself. Therefore, cheaters cannot arrange successful migrations and cannot violate the spatial structure mechanism. Hence, they encounter local extinction at their patches.

**Figure 2:**
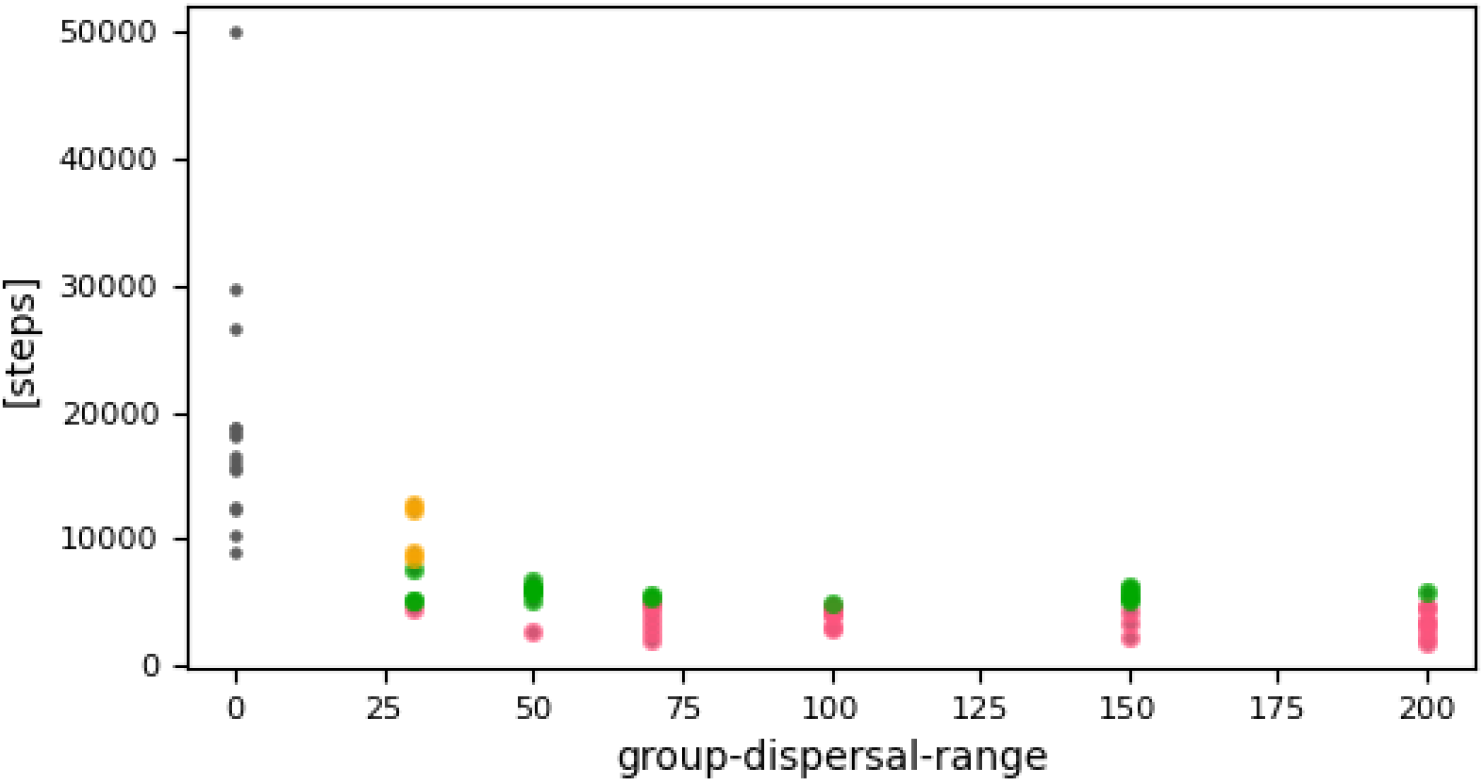
The runs that finished in favor of cheaters: (Dark yellow), above 8220 steps. (Light green), from 5000 to 8220 steps. (Pink), blew 5000 steps. The runs that finished in favor of cooperators: (Gray) All of them from 9000 to 30000 except one run persisted until the predefined stop limit of experiments 50000, with Three cheating agents as a final frequency.

**Figure 3:**
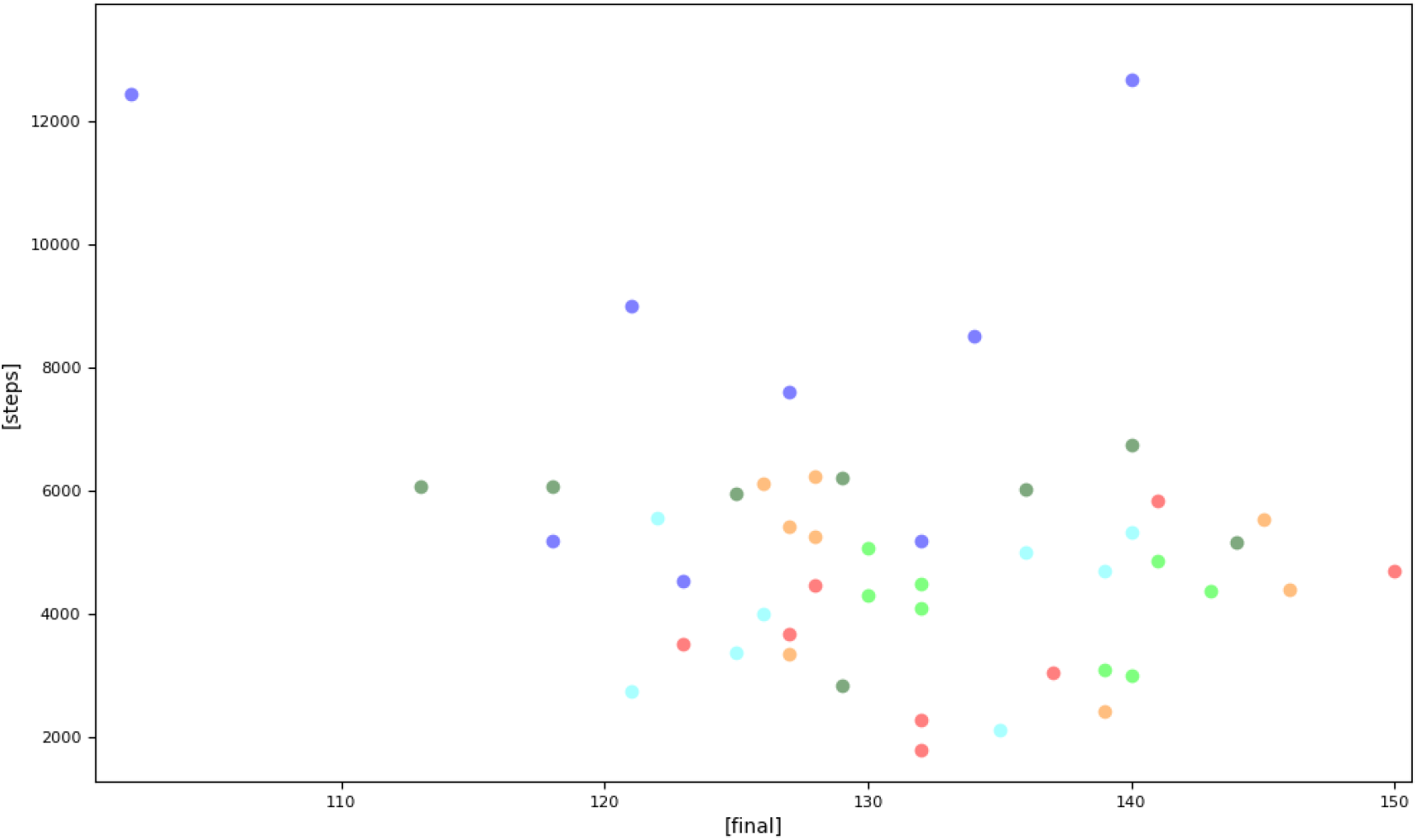
Group dispersal range = 30 (blue), 50 (dark green), 70 (sky blue), 100 (light green), 150 (orange), 200 (red). Cheaters did out-compete cooperators in all of these runs. However, the extinction of cooperators is likely to be done more quickly, with fewer steps in the higher group dispersal ranges.

The results agree with the intuitive predictions that cheaters could thrive, violate the spatial structure mechanism, and dominate the whole meta-population as long as they could cooperate to decrease the dispersal costs.

### 3.2. The experiments of the second model

The default values of the variables: Mutation rate = 1%. The kind of Punishment is suspended harvest once. Carrying capacity = 100. Number Agents = 250, (started number). Costs perception = 0.5. Growth rate = 0.3. Costs-punishment = 0.8. Percent-Punishers = 20%, (started ratio). Harvest-sustainable = 7. Percent-Sustainables = 99%, (most of the population consists of cooperators in the beginning). Living-costs = 4. Death-rate = 1. Perception-accuracy% = 99%, 90%, 70%, 60%, 50%, 40%, and 30%. Harvest-greedy = 15, 14, 13,12,11,10, and 9.

They are 44 runs of 7 experiments, all runs were 15000 ticks (iterations) and began with a 99% percent frequency of cooperators, and then they all finished up with greedy non-punishing taking over the population, the frequency of greedy non-punishing was above 90% in the final steps in all runs Fig. 4. And above 80% as a mean of all steps Fig. 5. I excluded the percent 100% accuracy, as it seems to me there is no such perfect monitoring case in nature. I began with 99% accuracy and then degraded to reach 30%, parallel to similar degradation in the greedy harvest amount from 15 to 9, Table. 1.

**Table.1:**
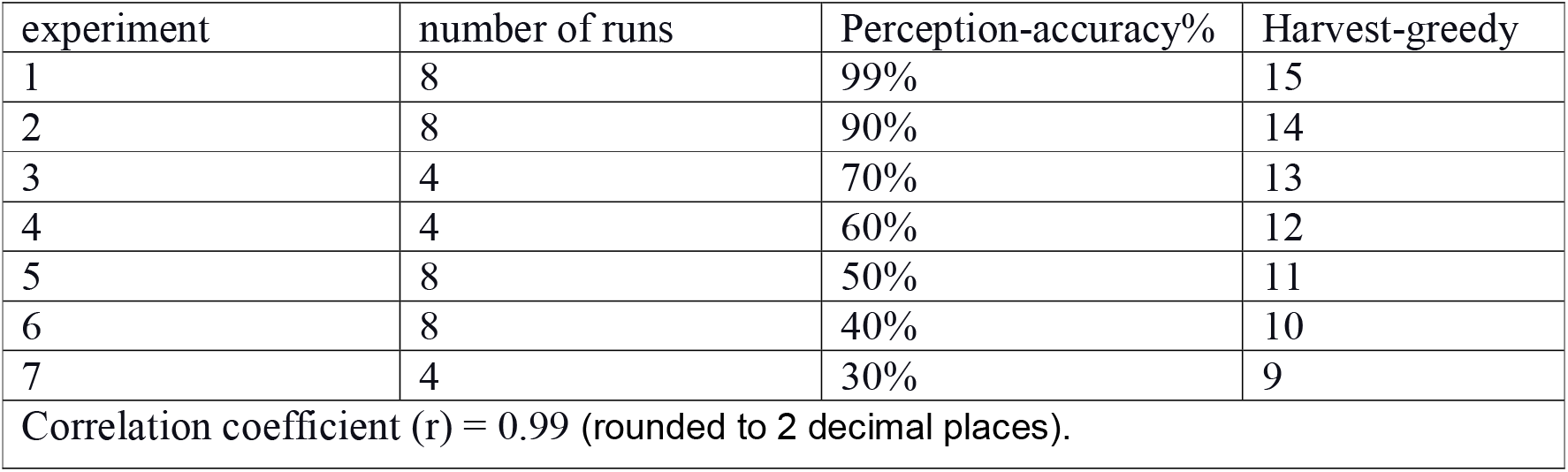

**Figure 4:**
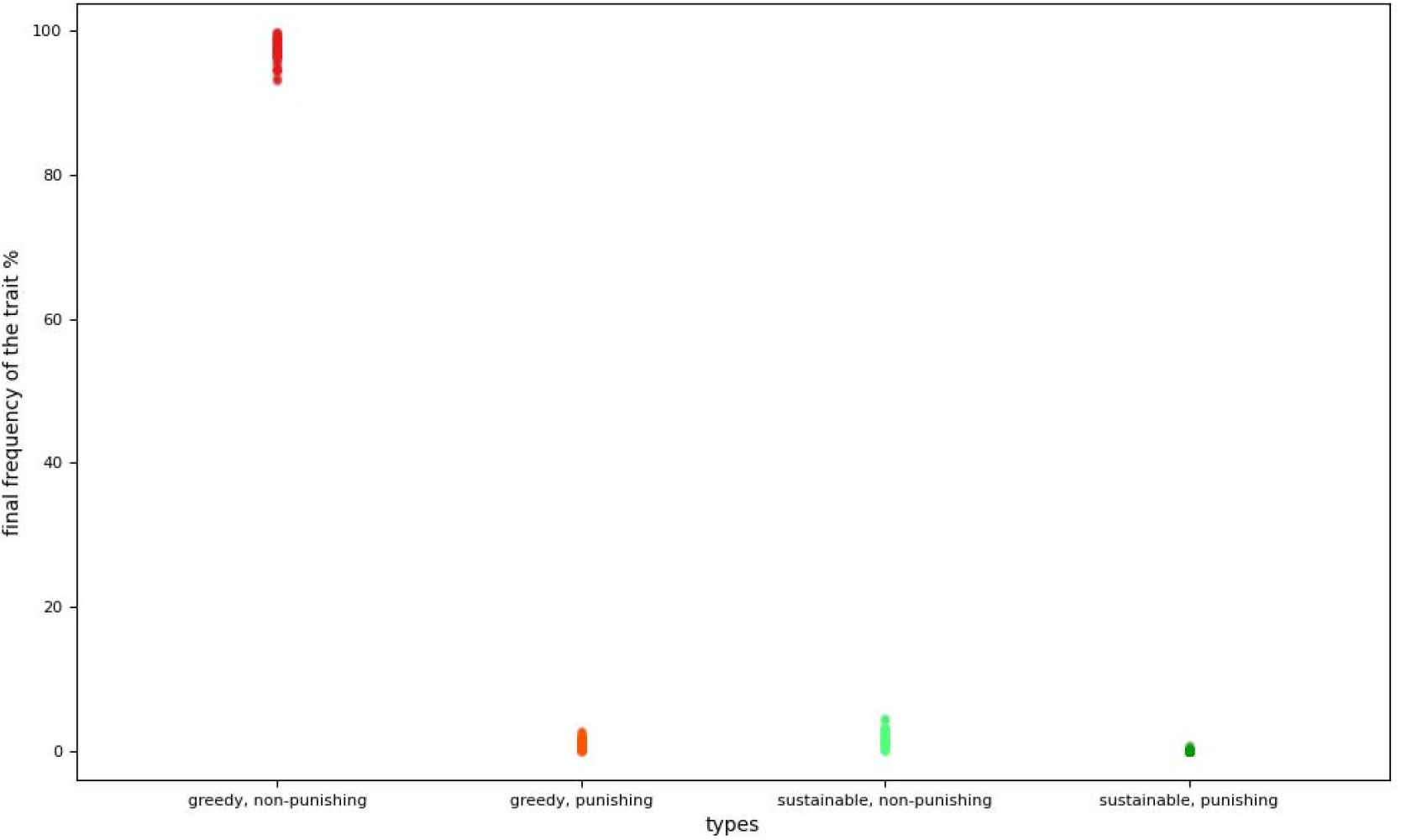
The four agent colors and types: 1) Red: greedy, non-punishing. 2) Orange: greedy, punishing. 3) Turquoise: sustainable, non-punishing. 4) Green: sustainable, punishing. The frequency of greedy non-punishing was above 90% in the final steps in all runs.

**Figure 5:**
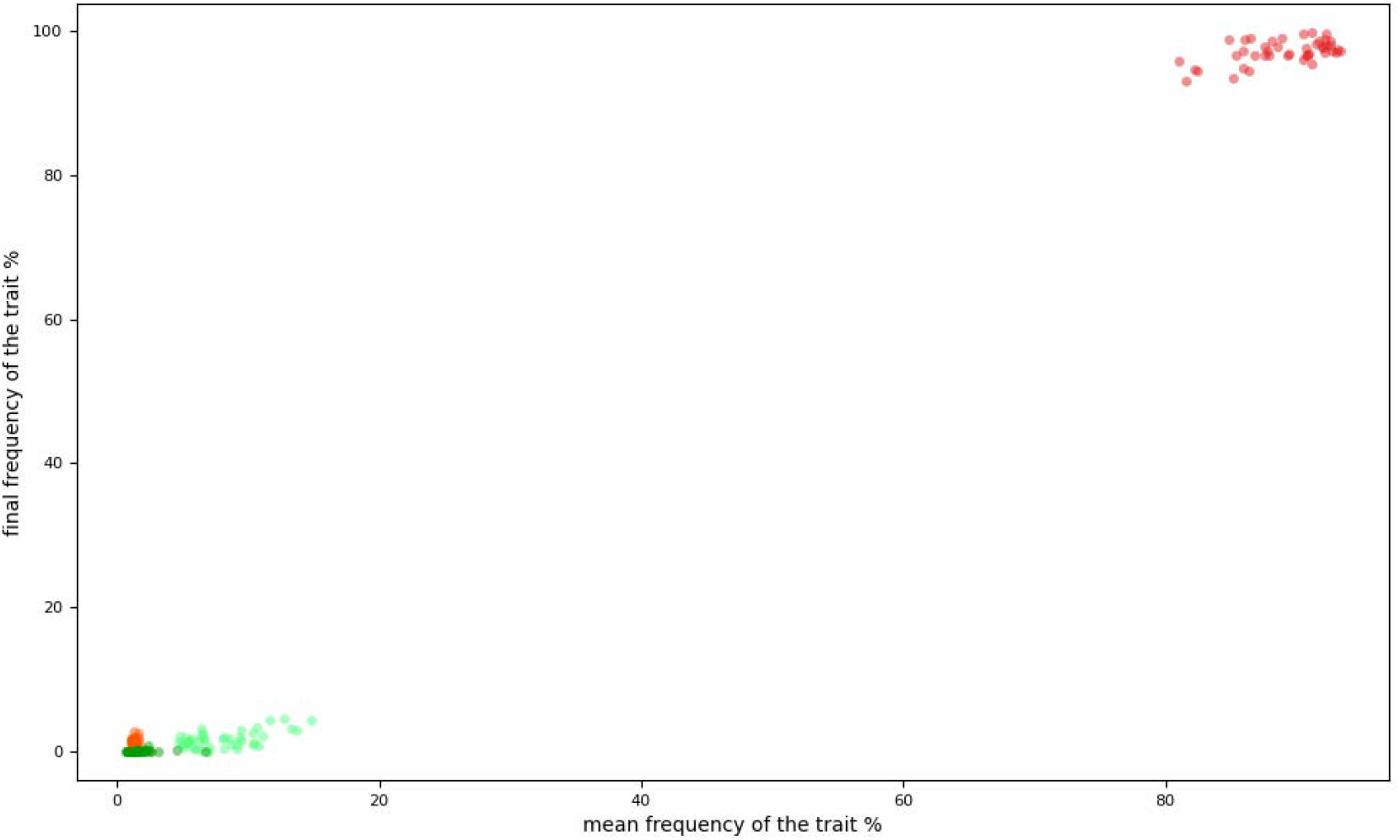
Combination between the frequency of The four agent types in the final and mean steps: greedy non-punishing (red) was above 90% in the final steps in all runs. And above 80% as a mean of all steps, also in all runs.

The results demonstrate a strong positive correlation (with correlation coefficient (r) = 0.99) between the values of the two variables (harvest greedy and perception accuracy). Which allow cheater dominance. Fig 6. Table. 1. Changing these values without considering this positive correlation prevents the dominance of cheaters (unpublished results). The dominance of cheaters means that they violated kin selection and punishment mechanisms.

**Figure 6:**
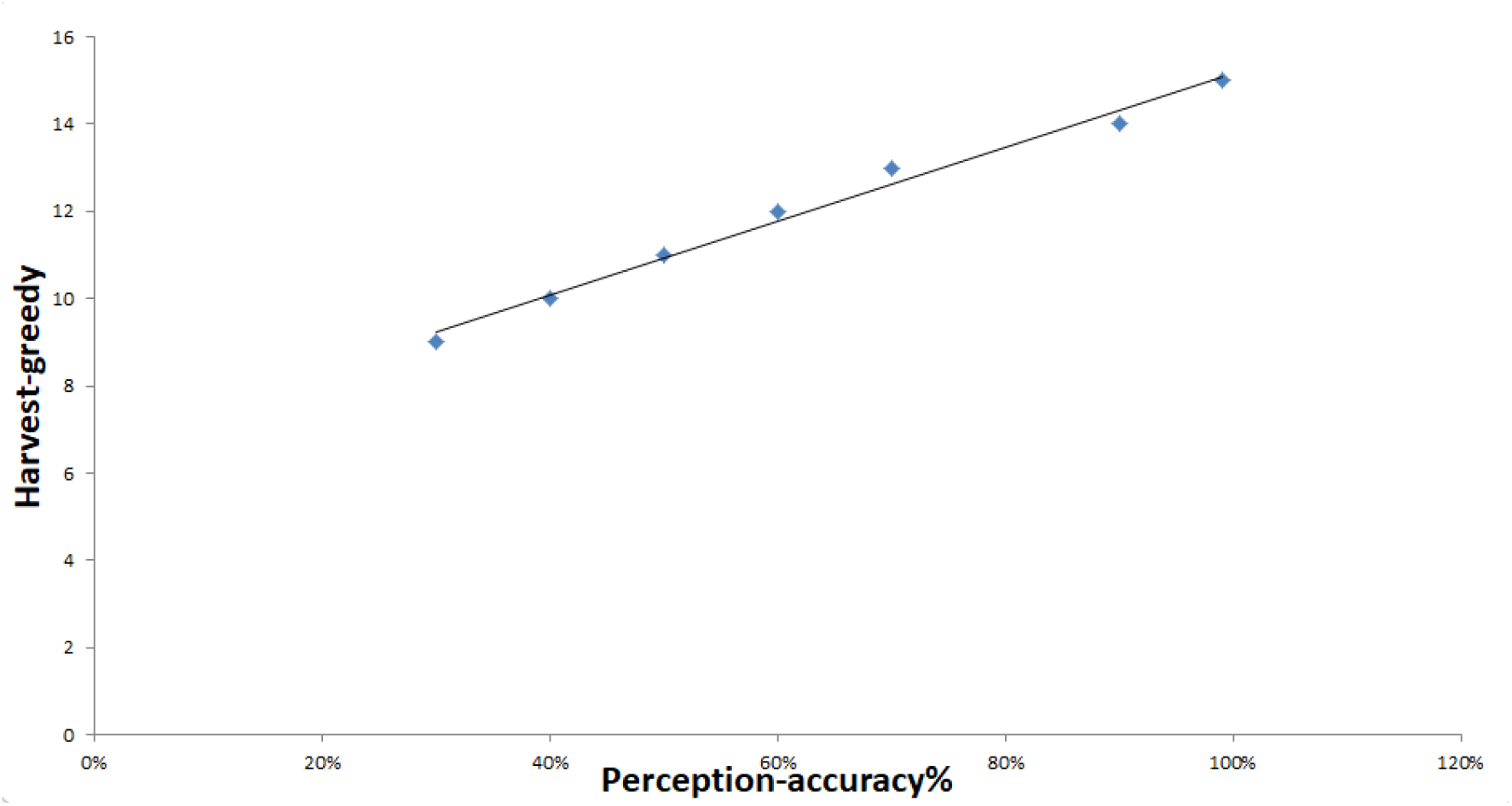
A strong positive correlation (with correlation coefficient (r) = 0.99) between the values of (harvest greedy and perception accuracy), which allow for cheater dominance.

## 4. Discussion

The conditional defector strategies can violate the most crucial supporting mechanisms of cooperation, such as spatial structure, kin selection, and punishment. This surprising outcome may cue us to rethink the evolution of cooperation as it illustrates that maintaining cooperation may be more difficult than previously thought. The two forms that I presented in this paper are general, as they have a broad range of applications and are simple. The zero-determinant (ZD) extortion strategy is also a conditional defector strategy if it has a tag-based decision to cooperate with relatives who adopt the same (ZD) extortion strategy but cheat otherwise. At that time, it could be stable and win the game against the opponent’s strategies, (Adami & Hintze 2013). In addition, when selfish strategies could modulate benefits and costs, they can outcompete tit for tat and generous strategies, (Stewart & Plotkin 2014). On the other hand, cheaters who can increase their dispersal rate without decreasing the dispersal costs often cannot achieve triumph, not drive the cooperators (wild type) to go extinct, or even harm themselves if the benefits of exploitation do not offset the costs of dispersal. For instance, the social parasite of *P. punctatus* ants is a wingless cheater queen. Although it has a high dispersal rate, it has costly migration on foot for long distances. Therefore, the colonies persisted for a long time instead of the supposed rapid collapse of the whole population, (Dobata et al. 2011).

The findings of the present paper suggest a potential therapy application. The conditional defectors can be used as suicidal agents to drive the population of pathogens into the self-destruction process. From an evolutionary perspective, tumors or microbes are considered populations consisting of cooperating cells that struggle for survival by adopting many collective costly actions to produce the intrinsic common resources, (Axelrod et al. 2006; Celiker & Gore 2013; West et al. 2007). Yet. Conditional defectors can violate the crucial mechanisms that support cooperation. Thereby, outcompeting the cooperators. The cheaters also would go extinct after cooperators because they cannot do the necessary collective actions. Undoubtedly, the production of the common resources or the public good I meant is not independent of cooperators like the two models in the present paper. Instead, its production ought to rely on cooperators. For example, the essential excretions of microbe deplete after the cooperator’s extinction. Cheaters can drive the whole population to go extinct; it is a well-established evolutionary prediction. This robust outcome appears in many theoretical and empirical studies and is known as the tragedy of the commons or evolutionary suicide, (Hardin 1968; Parvinen 2005; Rankin & LópezLSepulcre 2005). This phenomenon can occur if free riders have a fitness advantage over cooperators (wild-type) in an environment set by the cooperators. Creating the evolutionary suicide within the pathogen populations would mean the end of infections, or even endemics as cheaters are not static chemical substances but infectiously transmissible organisms.

It is not the first time someone suggests using cheaters in attacking pathogens as cooperator populations. For instance: (Brown et al. 2009). Suggested trojan horse therapy to reduce the virulence of pathogens or release beneficial medical substances inside its colonies.

(Notton et al. 2014). suggested the therapeutic Interfering Particles (TIPs) or hijacker therapy. It is a therapeutic utilize for the defective interfering particles (DIPs) that are molecular parasites of viruses or incomplete RNA particles lacking essential packaging elements. It is believed that it could defeat HIV and other viruses (like SARS-CoV-2). Moreover, (DIPs) are transmissible antivirals that can transfer from one person to another until ending the endemic in infected areas like sub-Saharan Africa, (Rast et al. 2016).

(Archetti 2013). Suggested autologous therapy. It aims to increase the diffusion range of the growth factors that the tumor is excreting. Hence, this could increase the tumor’s vulnerability to exploitation,(Archetti & Pienta 2019).

(Domingo-Calap et al. 2019). Manipulated a defector strain of *Vesicular stomatitis* virus called Δ51. And it doesn’t excrete a costly enzyme that suppressant the interferon. It could defeat the wild type. Then leads to the tragedy of the commons.

Other treatments and descriptive game-theoretic models of cancer are reviewed here,(Wölfl et al. 2021).

So far, many previous papers have suggested closely related ideas. But the defense mechanisms of cooperators were always a huge obstacle. I think now conditional defector strategies can surpass these obstacles.

## 5. Data and Software Availability

https://www.comses.net/codebases/8437728b-3e1f-46f3-80be-65f4f5909d81/releases/1.0.0/

## 6. Acknowledgment

I express thanks to my mentor Dervis Can Vural, Associate Professor. Physics Department, the University of Notre Dame, for his brilliant expert advice and encouragement. My colleague Mr. Karim Soliman for his extraordinary support and too long discussions on the construction of this study.

